# Evaluating Performance of Drug Repurposing Technologies

**DOI:** 10.1101/2020.12.03.410274

**Authors:** James Schuler, Zackary Falls, William Mangione, Matthew L. Hudson, Liana Bruggemann, Ram Samudrala

## Abstract

Drug repurposing technologies are growing in number and maturing. However, comparison to each other and to reality is hindered due to lack of consensus with respect to performance evaluation. Such comparability is necessary to determine scientific merit and to ensure that only meaningful predictions from repurposing technologies carry through to further validation and eventual patient use. Here, we review and compare performance evaluation measures for these technologies using version 2 of our shotgun repurposing Computational Analysis of Novel Drug Opportunities (CANDO) platform to illustrate their benefits, drawbacks, and limitations. Understanding and using different performance evaluation metrics ensures robust cross platform comparability, enabling us to continuously strive towards optimal repurposing by decreasing time and cost of drug discovery and development.

## 1 Introduction

Drug repurposing technologies allow us to predict new uses for previously approved drugs [1]. While the drug discovery process normally takes years of work and costs billions of dollars, drug repurposing can lower barriers to entry of a drug to the market [2,3]. The ultimate goal of repurposing research is to decrease time and cost of drug discovery and development by accurately predicting clinical utility and using the predictions to improve health and quality of life. Successful instances of drug repurposing have been based on anecdotal evidence, *in vitro* and *in vivo* screening, and discovery of serendipitous positive effects in clinical trials or analysis of patient health records post-market [4]. Drug repurposing technologies aim to make this process systematic and skip intermediate steps (Figure 1).

**Figure 1:**
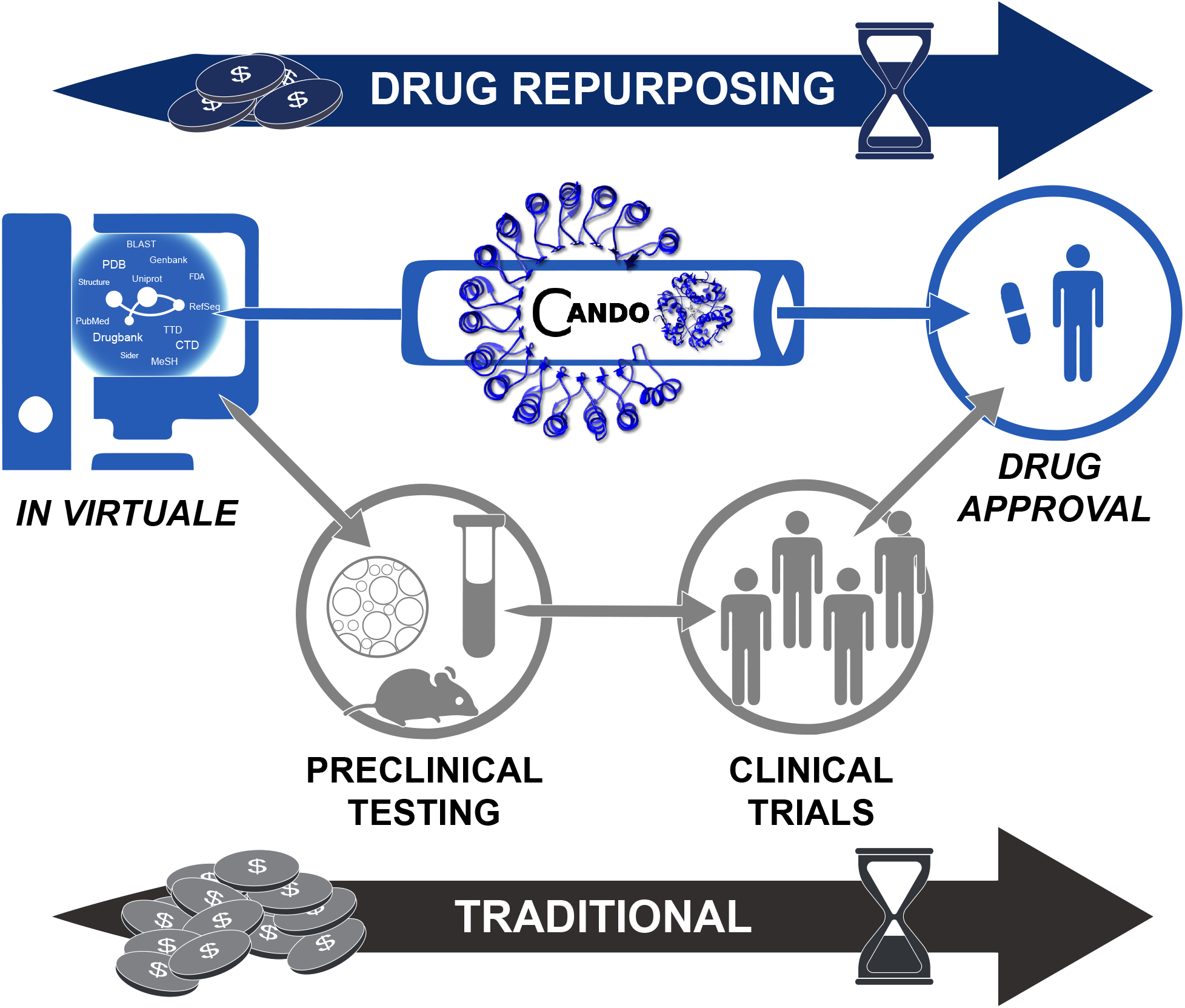
The relationship of drug repurposing technologies to traditional approaches. Traditional drug discovery and development is time consuming and costly, moving from preclinical research (basic computational methods, intensive *in vitro* and/or *in vivo* screening) to testing in clinical trials and eventual drug approval. The ideal scenario is a clinician/physician utilizing results of drug repurposing technologies (such as CANDO) directly, prescribing medications with high confidence to treat numerous indications, thereby saving time, cost, and improving patient outcomes. In this future guided by using the best evaluation metrics, repurposing technologies have high comparability and fidelity to reality.

Specific goals of drug repurposing differ based on their eventual utility. For a pharmaceutical company, a specific goal may be to find a single blockbuster drug which changes the default treatment of a condition shared by millions, for example, hypertension [5]. A basic science drug repurposing approach at an academic institution may focus more on public benefit and less on monetary outcomes, for instance, to find a treatment for an orphan or rare disease [6]. Less defined end goals (high risk) enable researchers to systematically disrupt the entire drug discovery and development process (high reward). Drug repurposing technologies such as our Computational Analysis of Novel Drug Opportunities (CANDO) platform [7–17] may be used to systematically predict the relative efficacy of every drug in its comprehensive library to treat every disease/indication, minimizing risks and amplifying rewards. In conjunction with mechanistic basic science analyses, these platforms may be used to better understand the science of drug behavior and thereby model reality with greater fidelity.

Distinctions between non-computational and computational drug repurposing technologies are blurring. Experiments designated as computational may rely on data collected in a bench environment such as protein-ligand binding energy measurement and gene expression studies [18]. Bench experiments may have used computational tools, such as homology modeling or molecular docking software [19], and computational studies are often externally validated or supplemented by *in vitro* and *in vivo* laboratory experiments [20]. This review and associated analyses focus largely on performance evaluation of computational technologies; however the metrics discussed herein are applicable broadly to drug repurposing, i.e., any technology that generates novel therapeutic repurposing candidates by benchmarking drug-indication association predictions.

Most previous reviews of drug repurposing technologies have focused on methods development [3,21–35], with only a few providing cursory analysis of evaluation of those methods [29, 35, 36]. Brown and Patel have previously reported a review of “validation” strategies for computational drug repurposing [37], broadly categorizing various evaluation metrics into “(1) validation with a single example or case study of a single disease area, (2) sensitivity-based validation only and (3) both sensitivity- and specificity-based validation” [37]. Shahreza et al., in reviewing network-based approaches to drug repurposing, mention various evaluation criteria used in different studies and provide mathematical aspects of the relationship between some of them [30].

Here we augment and enhance their foundational reporting by describing, reviewing, and analysing metrics for evaluating performance of drug repurposing technologies. We highlight uses of metrics borrowed from the realms of virtual screening/target prediction and information retrieval, and report results of their integration into CANDO. Through the use of better evaluation metrics, we aim to make drug repurposing science more rigorous and comparable. This study will help enable proper evaluation of drug repurposing technologies, and ultimately guide the field to bring about real changes in the armamentarium of medicine to alleviate disease burden.

### 1.1 Computational Analysis of Novel Drug Opportunities (CANDO)

We developed and deployed the CANDO platform to model the relationships between every disease/indication and every human use drug/compound [7–17]. Built upon the premise of polypharmacology and multitargeting, at the core of CANDO is the ability to infer similarity of compound/drug behavior. Canonically, we use molecular docking protocols to evaluate the interaction between large libraries of drugs/compounds and protein structures. We then construct a compound-proteome interaction signature to characterize and quantify their behavior. Based on the similarity of these interaction signatures, we rank every drug/compound relative to every other. We hypothesize that drugs/compounds with similar interaction signatures may be repurposed for the same indication(s).

Since the development and application of CANDO version 1 [7–10], we have continued to enhance our platform by analyzing the effect of protein subsets on drug behavior, implementing heterogeneous measures of drug/compound similarity, using multiple molecular docking software packages to evaluate interactions, and refining non-similarity based approaches for drug repurposing in situations where there there is no approved drug for a disease/indication [11–17,38].

### 1.2 Version 2 (v2)

Version 2 of the CANDO platform (v2) described here, implementing updated drug/compound and protein structure libraries, indication lists, drug-indication mappings, interaction scoring protocols, benchmarking and evaluation metrics, along with data fusion of multiple pipelines mixing and matching between these choices, is used as a template for the rigorous evaluation of the performance of drug repurposing technologies. The core tenets remain the same as in CANDO v1; however, the updated data and evaluation metrics enable us to better determine the correctness of those predictions with greater cross-platform comparability, as well as agreement with preclinical and clinical validation experiments.

#### 1.2.1 Curation of drug/compound and protein structure libraries

The v2 compound library contains 2,162 US FDA approved drugs extracted from DrugBank 5.0 [39]. We have also created a larger library of 8,752 compounds from DrugBank containing both approved drugs and experimental/investigational compounds. The updated protein library contains a nonredundant set of 14,606 solved structures compiled from the Protein Data Bank (PDB) [40]. Supplementing the approved drugs with compounds that are in the final stages of the drug development process helps to expand the repurposing capabilities of the platform. The updated protein structure library comprises of individual chains from numerous proteomes with an equitable distribution of folds and binding sites, allowing for greater coverage due to homology while reducing bias.

#### 1.2.2 Interaction scoring

The default pipeline in CANDO v2 uses an enhanced version of our previous bioinformatic docking protocol for interaction scoring [8]. We altered the previous protocol by using extended connectivity fingerprints from RDKit [41] instead of FP2 fingerprints from OpenBabel [42] as our cheminformatic approach for drug/compound molecular fingerprinting. The other major modification is using COACH to predict protein structure binding sites [43]. COACH leverages the results from multiple binding site prediction software suites, including COFACTOR which we previously used exclusively. The use of the updated fingerprints and COACH results in higher fidelity to reality for the resulting interaction scores from the bioanalytics-based docking protocol, as evaluated by comparing to observed compound-protein interaction binding constants obtained from PDBBind [12].

#### 1.2.3 Drug/compound characterization, benchmarking, evaluation metrics, and performance

The default pipeline in CANDO v2 generates drug-proteome interaction signatures by calculating scores between all 2,162 drugs and all 14,606 proteins using the bioanalytics-based docking protocol. Every drug is characterized by a unique vector of interaction scores. We measure the pairwise similarity/distance between the interaction signatures from each of the 2,162 drugs to every other. We evaluated a variety of similarity measures between a pair of interaction signatures and found success using the root mean squared deviation (RMSD), which is the default measure, and the cosine distance [44]. We then sort drugs relative to one another according to their similarity, i.e., those with the greatest similarity to each other are ranked highest.

Each drug is associated with one or more approved indications, comprising our standard to which we compare our rankings (also known as a “gold standard” or “ground truth”). These associations, or drug-indication mapping, is derived from curating the Comparative Toxicogenomics Database (CTD) [45], which uses medical subject headings (MeSH) to label indications [46]. This yields 18,709 associations for the 2,162 drugs covering a total of 2,178 indications. The default benchmarking protocol implements an in-house leave one out procedure to accurately identify related compounds approved for the same indication [8, 15]. For every indication associated with a drug, we calculate the ranks of other drugs associated with that same indication, and whether any positive hit occurs within certain cutoffs, such as top10, top25, etc. representing the top 10 and 25 most similar drugs. For each indication, we calculate the percent of associated drugs which achieve a hit in that cutoff. We next calculate the mean of all per-indication accuracies to give an overall evaluation of the platform, referred to as the average indication accuracy (AIA) [8,15].

For drug repurposing, our small molecule library is limited to the 2,162 approved drugs but CANDO is capable of analysing compounds that are not yet approved in a similar fashion. Similar drugs/compounds not associated with the same indication are hypothesized to be novel repurposed therapies to be validated via external preclinical and clinical studies.

Consider our results for the indication melanoma (MeSH ID: D008545), which is associated with a curated list of 58 drugs from the CTD. CANDO predicts 23 of those 58 drugs to have another associated with melanoma within their respective top 10 most similar proteomic interaction signatures. Therefore, the top10 indication accuracy for melanoma is 23/58 × 100 = 39.6%. We repeat this process for every one of the 2,178 indications to generate the AIA.

For version 1 (v1) of CANDO, we achieved a top10 AIA of 11.8%, compared to a reported random control of 0.2% [8]. Using improved bio- and cheminformatic tools and armed with a better understanding of random controls, theoretically modeled using a hypergeometric distribution and empirically measured using uniformly random drug-drug similarity data, version 1.5 of CANDO achieved a average indication accuracy of 12.8% at the top 10 cutoff against a random control of 2.2% [12].

The average indication accuracy is not used by others in the field of drug repurposing technologies. Thus, while we use it as a metric for internal comparison (i.e., between individual CANDO pipelines and versions), the cross-platform applicability is low. We thus researched other methods of assessing performance which are now implemented in and applied to CANDO.

### 1.3 Classification, ranking, metrics, and integration into CANDO

Experiments using drug repurposing technologies may return results as a classification or ranking. In classification, compounds are associated with indications in a binary fashion based on some criteria, whereas in a learning to rank experiment, entities are ranked relative to one another in order of some score. Best pairings, designated by a specific rank/classification cutoff/threshold are reported as putative therapeutics. A ranking result can be thought of as classification by using the cutoff as a threshold for the ranks, and declaring items on one side of the threshold as positive samples, and those on the other side as negative samples. The inverse of modeling classification as a ranking problem is true as well. If a specific cutoff is used for calculation of performance, it must be reported, and reporting values at all possible ranking and classification cutoffs/thresholds will allow for greater comparability.

An inherent limitation of drug repurposing technologies and nearly all metrics we use to report on their goodness is the forced dichotomization of results, where those drug-indication associations ranked/classified better than the cutoff/threshold are labeled “positive” and those worse are “negative”, with no nuance or allowance for real-world considerations such as first- and second-line therapies. This lowers the fidelity of all drug repurposing models to reality. Figure 2 illustrates these issues in a *mock example* of predicted therapies to treat type 2 diabetes by a drug repurposing technology.

**Figure 2:**
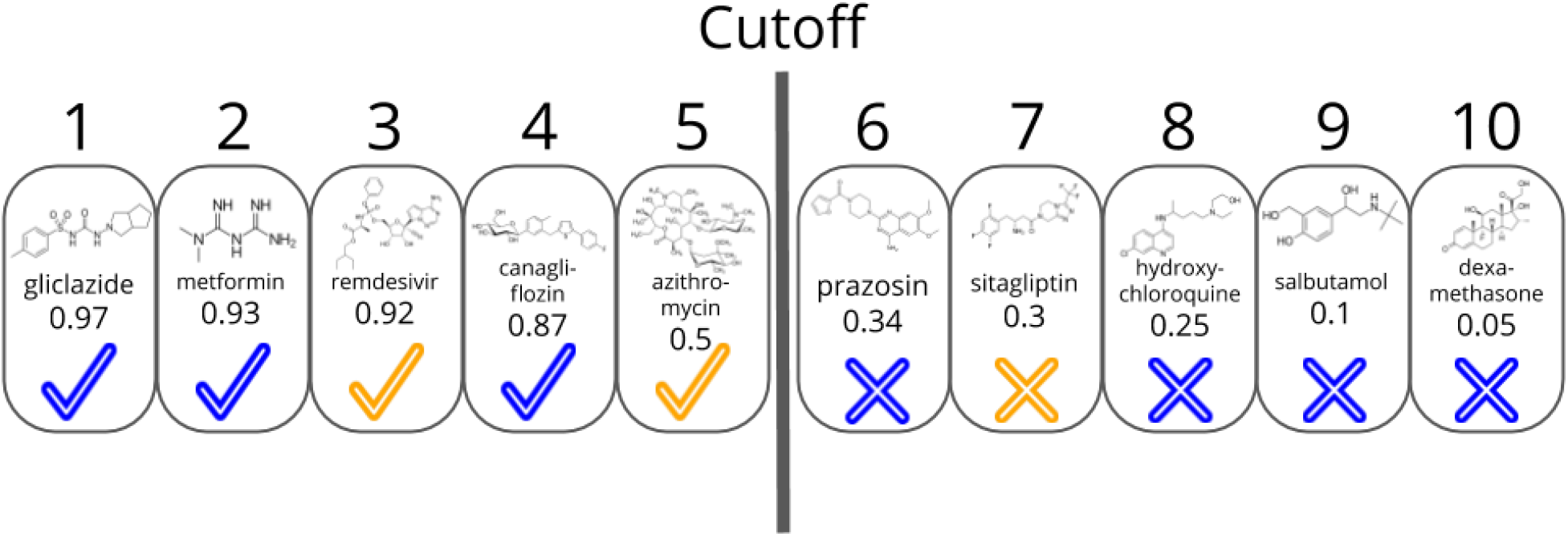
Mock example of novel treatment prediction for type 2 diabetes by a drug repurposing technology. We illustrate an arbitrary methodology which ranks 10 drugs to treat type 2 diabetes, with rank one (number above boxes) being the most likely efficacious and rank ten being the least likely. These ranks are based on *mock scores* given below the name of each drug. Classification of compounds with a score ≥ 0.5 are the same as those ranked in positions one through five, i.e. the results of a classification and ranking schemes are interchangeable. Drugs which have known treatment associations with type 2 diabetes are marked with a blue check or cross, and those with unknown associations are marked correspondingly in orange. Those drugs classified/ranked better than the cutoff are the positive results, whereas those worse are the negative results. As there are three drugs with a known association to type 2 diabetes (blue check) and two with no known association (orange check) ranked better than or equal to rank five, there are three true positives and two false positives. Analogously, there are four true negatives (blue cross) and one false negative (orange cross). If this were not a mock example, the false positive results would be the repurposing candidates for the given indication. We also note the notion of “true negative” can be misleading, as the vast majority of such associations are of undetermined classification, having never been rigorously scientifically studied. This lack of negative data in comparison standards is a limitation in the evaluation of drug repurposing technologies. This is a mock example of a single ranking among hundreds or thousands in a comprehensive drug repurposing platform. Metrics utilizing a single ranking are averaged over many rankings to produce values which describe the correctness of a drug repurposing technology.

Drug repurposing technologies use a multitude of different metrics to evaluate performance of the resulting ranking or classification. These are based on delineation of results into true positives, false negatives, true negatives, and false positives relative to some standard. Notable commonly used metrics include sensitivity (true positive rate, recall), specificity (true negative rate), false discovery rate, false positive rate precision (positive predictive value), area under the receiver operating characteristic curve (ROC and AUROC), precision, precision-recall curves, and area under precision-recall curves, F1-score, and Matthews correlation coefficient (MCC, Supplementary Material).

Results of a drug repurposing experiment composed of a ranking of drug candidates are similar to those results obtained in virtual screening and target prediction experiments, but the standard of comparison is different: known drug characteristics or drug-target associations (binding interactions) as opposed to known drug-indication associations. Drug repurposing as a field is not one-to-one with drug target prediction [37], but a drug target prediction experiment can be part of a repurposing experiment. Despite this demarcation, metrics used in virtual screening/target prediction highlighting the “early recognition problem” may be quite useful to evaluate drug repurposing [47,48]. In both instances the goal is to prioritize ranking “active” candidates (known drug-target or drug-indication associations) at the top. Metrics that consider the early recognition problem properly include the enrichment factor (EF, Supplementary Material), robust initial enhancement (RIE, Supplementary Material) [49], and the Boltzmann-enhanced discrimination of receiver operating characteristic (BEDROC) [47].

In addition to being similar to virtual screening, the results of drug repurposing technologies are highly analogous to information retrieval. Information retrieval tools such as web search engines may be evaluated in their ability to accurately return a website link that is desired and subsequently visited by the user based on some query string. In a drug repurposing experiment, this is akin to generating a list of active candidate drugs which may be a treatment for a given indication. Therefore, the goals of information retrieval and drug repurposing are similar, and correspondingly performance metrics that have been explored for information retrieval have value in drug repurposing evaluation. These include the mean reciprocal rank (MRR), precision-at-K (P@K), average precision (AP) and mean average precision (MAP, Supplementary Material), and [normalized] discounted cumulative gain ([N]DCG) [50].

We describe the utility and use of the above mentioned metrics in several computational drug repurposing experiments. *We report complete results of using every metric at all cutoffs for both versions 1.5 and v2 of CANDO in the Supplementary Material and at our website, and highlight specific results of interest in the following text and figures.* In general, most metrics have use for internal intra-platform comparisons but limited use for external inter-platform comparisons. Based on our analyses, we conclude that the metrics with the most utility relative to their cost are BEDROC and NDCG. Additionally, we have integrated many of these measures into the CANDO platform to facilitate internal and external comparability. Evaluation of CANDO using these newly integrated metrics has reaffirmed its utility for drug repurposing, while providing a foundational review of the advantages and limitations of each metric in the context of the libraries and standards used.

## 2 Measures of correctness/success

### 2.1 Mean Reciprocal Rank

Reciprocal rank is the inverse of the position of the first correctly retrieved active in a ranking scheme, or the best scoring active in a classification. The correctness/success is determined by matching the retrieved active to a known drug-indication association according to some standard. Mean Reciprocal Rank (MRR) is the average of the inverse rank of each first retrieved active, i.e., the first true positive [51]. While easy to calculate, this metric only uses the ranking of the first retrieved active. According to MRR, a drug repurposing experiment that ranks a single active correctly out of several performs just as well as another that ranks several correctly. An overall measure of correctness is difficult to discern from reporting of a single value; however, a possible way to evaluate distributions of performance across several experiments is provided in [51].

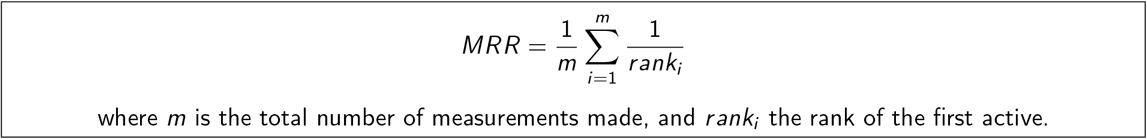

MRR is the least similar to the other metrics reviewed. It does have utility ranking putative drug candidates for an indication for which there is a single known association, i.e., there is only one active to compare against for evaluation of correctness/success, such as with some neglected and emerging indications. Our previous metric for evaluating performance in CANDO, AIA, is similar to MRR, in that for each drug-indication pair, we are primarily concerned with whether there is an active within a certain cutoff.

### 2.2 True Positive, True Negative, False Positive, False Negative

Given a classification or ranking of candidate drugs for repurposing, the positive samples retrieved are those deemed to be associated with the desired indication or correctly ranked within the specified cutoff. Negative samples are those classified as having no association with the desired indication or ranked outside the cutoff. Within these, there are true positive (TP), false positive (FP), true negative (TN), and false negative (FN) samples. True positive samples are correctly classified or ranked associations that are present in the standard, whereas false positives are incorrectly classified as positive or ranked within the cutoff, with the corresponding association not in the standard. False negatives are known associations found in the standard but classified incorrectly or ranked outside the cutoff, whereas true negatives are not present in the standard and correctly classified as such or ranked outside the cutoff.

### 2.3 Sensitivity and specificity

Sensitivity is the proportion of true positives that are correctly identified; specificity is the proportion of true negatives that are correctly identified:

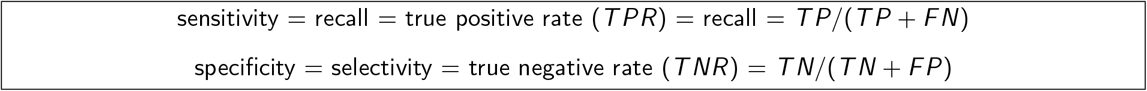

Lim et al. [52] used sensitivity to report drug-target prediction correctness using their REMAP platform. The authors did not directly quantify their drug repurposing predictions, but found corroborating examples of novel treatments in the literature. Donner et al. use recall as a metric for reporting and visualization of ranking perturbagens (chemical substances which change gene expression) [53], and Xuan et al. graphically show the recall at top cutoffs of rankings of drug-indication associations generated via their methods [54]. Wu and colleagues use sensitivity as one of their metrics of choice to evaluate the ability of their repurposing platform (MD-Miner) to identify active drugs among top ranked candidates for repurposing [55]. The reporting of sensitivity and specificity is usually not the focus in these studies. Instead, they are described and displayed as part of another metric, often the receiver operator characteristic curves and area under such curves discussed below. Figure 3 illustrates application of this metric to CANDO.

**Figure 3:**
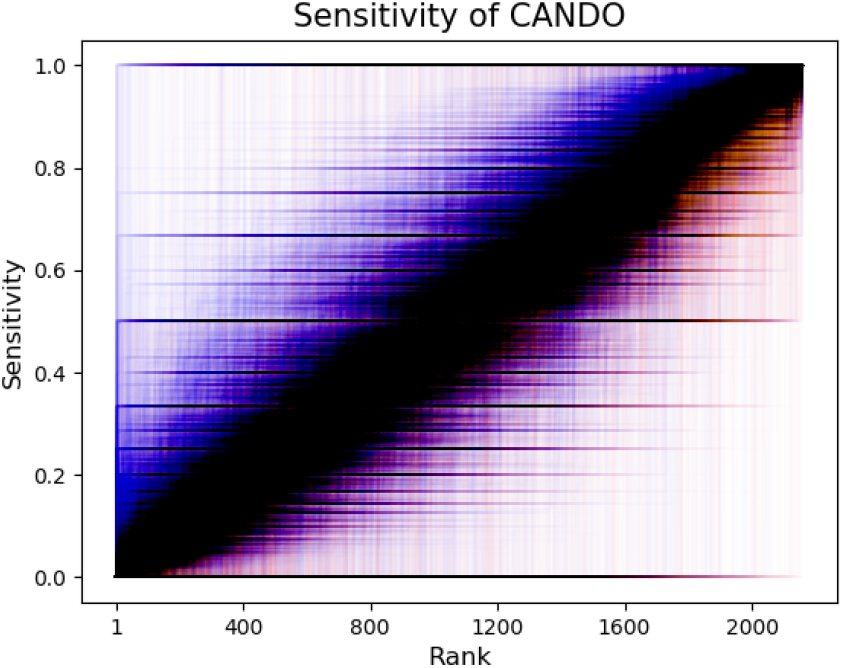
Evaluating CANDO performance using Sensitivity. The sensitivity values (vertical axis) of all drug-indication associations in CANDO v2 (blue) and a random control (red) are shown according to the rank (horizontal axis). Broadly, more drug-indication pairs score better at all ranks using the v2 pipeline relative to random, visually observed as more blue in the upper left half and purple in the lower right half above. The darkness of a point is directly proportional to how many lines pass through that point. This is the display of the results using a single metric for a single pipeline within a single platform compared to the same pipeline with random input data, highlighting difficulties in illustrating data of this type and size. More metrics, more pipelines, more platforms, and more data (including controls), will greatly increase illustration complexity.

### 2.4 False Discovery Rate and False Positive Rate

More true positives are classified and ranked correctly when quantifying and subsequently limiting the number of false positives. In drug repurposing, the false discovery rate (FDR) is typically used as a cutoff for further development of specific results and not as a standalone metric. Through consideration of drugs, inflammatory bowel disease (IBD) genes, and biological pathways, Grenier and Hu generated candidate treatments for IBD, using the FDR as a way to guide classification of putative therapeutics [56]. Sirota et al. use FDR to measure significance of drug-indication scores compared to random in their experiments comparing gene expression levels as drug signatures [18]. Hingorani et al. claim drug discovery projects fail because of their excessive FDR [57]; in addition, they calculate the probability of repurposing success based on several assumptions. A utility of drug repurposing technologies is to reduce the FDR, but current methods may easily inflate the number of false discoveries by generating excessive numbers of predictions [58]. Lim et al. use a confidence weight to quantify uncertainties in predictions to reduce false positives [59]. In a similar way to the FDR, the false positive rate (FPR) is generally used as an integral part of another metric.

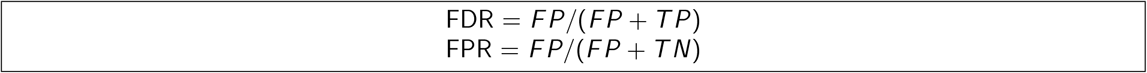

### 2.5 Receiver Operating Characteristic Curve and Area Under the ROC Curve

Receiver operating characteristic (ROC) curves are graphs where each point is the representation of a binary classification performance measured using the true and false positive rates. The points along an ROC curve are discrete, but often shown as continuous lines that are obtained by varying the cutoff for classification or rank value and calculating the corresponding TPR and FPR. One of the more popular methods for assessing and reporting performance of drug repurposing technologies is the area under the ROC curve (AUC, AUROC). A singular value, the AUROC is calculated using the trapezoid rule, or directly using the rank of the actives (drug-indication associations present in the standard) along with the ratio of actives and ratio of inactives (to be determined associations, or not present in the standard) in the entire drug library [47]. We have implemented the second approach in CANDO (Figure 4). A higher value is taken to be indicative of better performance, with a perfect classification obtaining an AUROC of 1.0, and 0.5 indicating random ranking/classification.

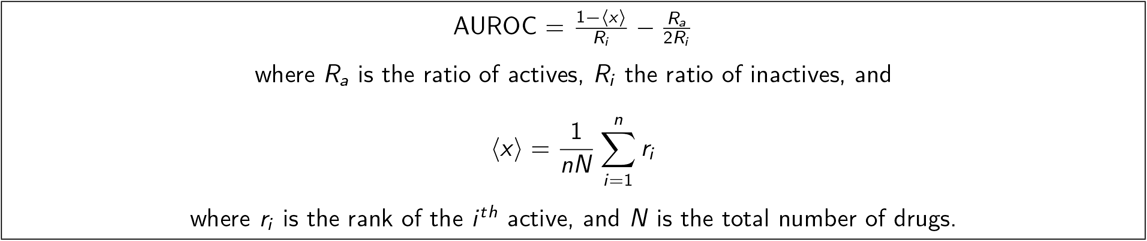

**Figure 4:**
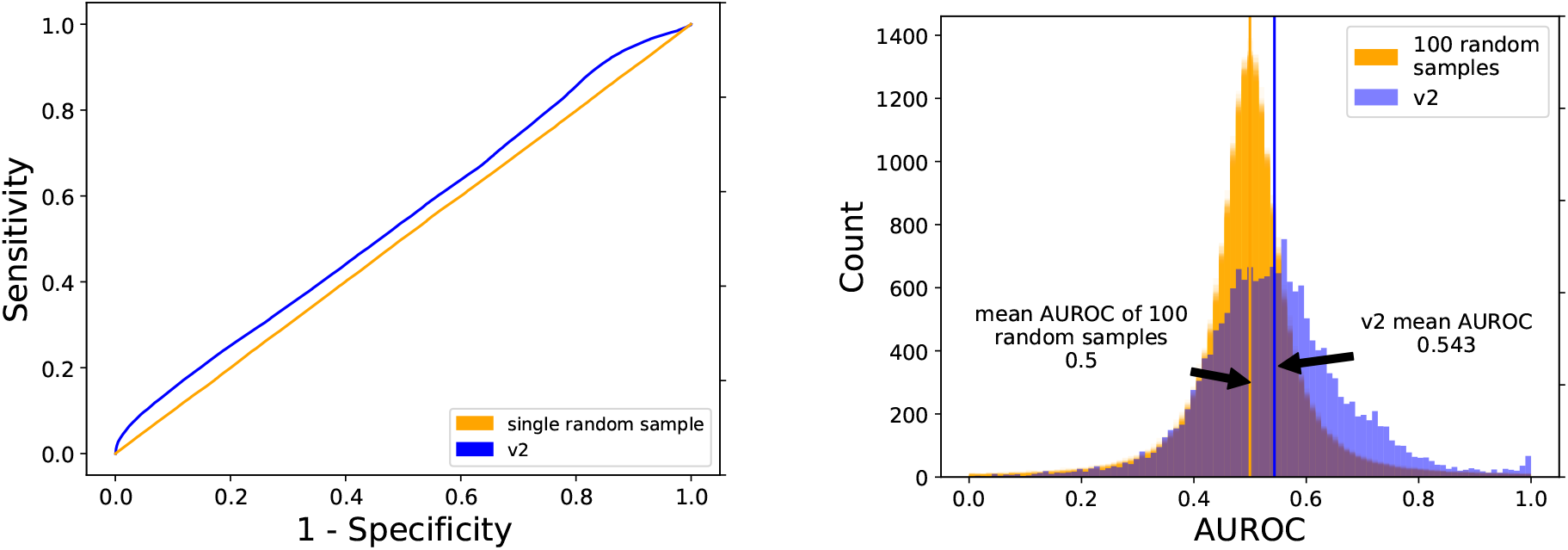
Evaluating CANDO performance using ROC and AUROC. The left panel shows the mean ROC curve across all drug-indication pairs for CANDO v2 (blue) and a single sample random control (orange). As with all ROC curves, the horizontal axis is 1 - specificity and the vertical axis is sensitivity. The empirical random matches what is mathematically expected by random (a straight line along the diagonal), i.e., both reflecting drugindication associations obtained by chance based on a uniform distribution. The right panel shows a histogram of AUROC scores for all drug-indication pairs predicted by v2 (blue) compared to those from 100 random runs (orange). The mean AUROC of all drug-indication pairs from v2 is 0.543, compared to a empirically derived random mean of 0.5 (again, matching theoretical expectation). The right shift of the v2 AUROC indicates an overall better performance relative to random controls. Both ROC and AUROC are useful for internal validation, but overall have limited utility in assessing different drug repurposing experiments or technologies, in part due to imbalances in known drug-indication associations and not emphasizing early recognition.

The drug repurposing project PREDICT uses AUROC as a main method of reporting goodness [60]. Moridi et al., report their own AUROC compared to that of PREDICT [61]. Nguyen et al. create a computational drug repurposing framework based on control system theory (DeCoST) to make novel treatment predictions for cancer [62]. Notably, Nguyen et al. include negative associations in their studies, such as drugs withdrawn from treatment or terminated clinical trials, which enhances the fidelity of their computational experiments to reality [62]. Due to different libraries and standards used, drug repurposing studies are not ideally analyzed through the use of the AUROC. As stated previously, the AUROC is highly dependent on the ratio of actives to inactives in a library; the DeCoST framework did overcome this dependency issue in part by creating a new, more balanced, library derived from drugs used by another group [63]. Emre Guney used AUROC as a metric, pointing out how data in a drug repurposing experiment may bias scientists toward conclusions that are not justified, and how single values of AUROC may not hold up to further cross-validation [64].

While AUROC may have low comparability between platforms, Lee et al. nonetheless compared results of their drug repurposing experiments across a diverse breadth of indication types, and showed better performance than those of others for the same indication types [65]. The mean AUROC of CANDO v2, calculated on a per indication basis, which itself is a corresponding average of grouped drug-indication pairs when there is more than one drug, is 0.520, with a median of 0.525 (interquartile range: 0.481 - 0.561).

The biggest shortcomings of the ROC/AUROC is the lack of early recognition and inability to handle imbalanced data. As illustrated with respect to virtual screening [47], the metric fails to enable comparison of drug repurposing for ranking actives at the top of an ordered list, which is the desired goal.

### 2.6 Precision and Precision-Recall, and Area Under the Precision-Recall Curve

Precision measures the relevance of a set of predictions:

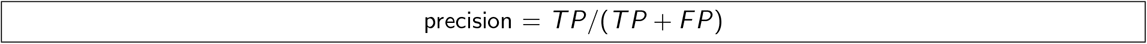

Yu et al. use established disease-gene-drug relationships to infer new drug-tissue-specific-disease relationships, and report precision as a standalone metric (as a score relative to the top percent of drug-indication pairs) [66]. Precision is often reported alongside recall, and one of the most commonly used metrics used to evaluate drug repurposing technologies is area under the precision-recall (PR) curve (AUPRC). Saito and Rehmsmeier provide strong evidence on the superiority of precision-recall compared to ROC when evaluating imbalanced data [67]. Imbalanced data is commonplace in drug-indication association standards used by repurposing technologies; i.e., the drugs in the standard are spread divergently across the indications, and vice versa.

PR curves may show a distinctive jagged edge pattern or appear finely interpolated, retaining maximum precision up to a particular recall [50]. Xuan et al. use both ROC and PR to compare their drug-indication association predictions to those made by others [54]. Peng et al. also use ROC and PR curves to demonstrate the internal validation of their network-based inferences about drug Anatomic Therapeutic Classification (ATC) codes [68]. McCusker use precision to evaluate their computational drug repurposing predictions to treat melanoma as “the percentage of returned candidates that have been validated experimentally or have been in a clinical trial versus all candidates returned” [69]. This metric is indeed precision, albeit based on a different type of standard, as they apply it to evaluate capturing literature instead of known associations.

Iwata et al. use AUPRC to assess internal performance of reconstructing known drug-indication associations made using supervised network inference, and compare their results to those obtained by others [70]. Khalid and Sezerman report AUPRC along with AUC and mean percentile rank to demonstrate the ability of their platform, which combines protein-protein interaction, pathway, protein binding site structural and disease similarity data, to capture known drug-indication associations [71]. With respect to cross-platform comparability, the authors applied their algorithm to evaluate performance using gold standard data available online for three other platforms and found that they obtain better AUC values using their methods but with data from other platforms [71].

### 2.7 Accuracy and F1-score

The term “accuracy” causes confusion since it is used to refer to performance generally in a colloquial sense, and also to a mathematically defined value in the context of binary classification. In a drug repurposing evaluation context, accuracy is the fraction of true positives and true negatives correctly classified:

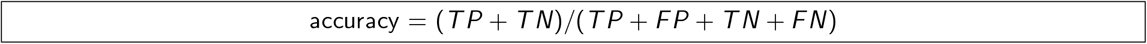

Accuracy is extremely influenced by the number of actives and inactives in a set and its utility is therefore limited. Accordingly, it is best used with balanced data, which is rare in drug repurposing technology standards. CANDO v2 obtains an average accuracy over all drug-indication pairs of 0.94. This high value is appealing at first, but is useless as it is identical to the mean average accuracy over all pairs for data collected over 100 random sample runs. This is due to our standard being greatly skewed in (correctly) representing the known drug-indication associations.

The F1-score (F-score, F-measure), widely used in machine learning applications, is the harmonic mean of precision and recall:

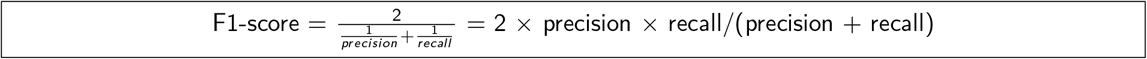

By focusing on small value outliers and mitigating the impact of large ones, the F1-score provides an intuitive measure of correctness when using uneven class sizes, unlike accuracy. Just as precision and recall are calculated at a certain cutoff, the ranks at which the measurement is made, or the score used for classification, should be reported along with the F1-score. Using CANDO v2, we obtain a mean F1-score calculated over all drug-indication pairs at the top 100 cutoff of 0.033, compared to the mean of 100 random samples at the same cutoff of 0.023.

Zhang et al. use AUC, precision, recall, and F1-score in evaluating their SLAMS algorithm [72]. Their calculation of recall is not directly related to the other measures [72], an indication of how different groups measure performance variably. Aliper et al. report results of their deep learning platform for drug repurposing based on transcriptomic data using accuracy and F1-score [73]. Specifically, the authors not only report averages, but also performance of their platform across three, five, and twelve specific therapeutic use categories according to the MeSH classification, and putative explanations for differences. McCusker et al. also use F1-score in evaluating performance of their probabilistic knowledge graph platform [69].

### 2.8 Boltzmann-Enhanced Discrimination of Receiver Operating Characteristic

The Boltzmann-Enhanced Discrimination of Receiver Operating Characteristic (BEDROC) metric merges early recognition with the area under the ROC curve [47]. More formally, the BEDROC metric evaluates the probability that an active ranked by the evaluated method will be found before any other that is derived from a hypothetical exponential probability distribution function with parameter *α*, where *αR_a_* ≪ 1 and *α* = 0. In this context, *R_a_* is the ratio of actives in the standard and 1/α is “understood as the fraction of the list where the weight is important” [47]. The authors state that BEDROC should be understood as assessing “virtual screening usefulness” as opposed to “improvement over random” (which is what the ROC does). In the context of drug repurposing, this may be interpreted as probability that a drug predicted to treat an indication is ranked better than a drug that is not.

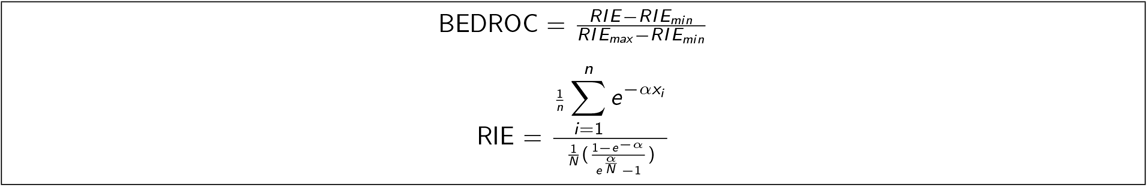

RIE is itself another metric known as robust initial enhancement. *RIE_min_* is the calculated RIE when all actives are at the bottom of a ranked list, *RIE_max_* when they are all ranked better than any inactives, *x_i_*, is the relative rank of the *i^th^* active, i.e., 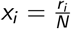, where *r_i_* is the rank of the active, *N* is the total number of drugs/compounds, *n* is the number of actives, and 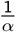 is “the fraction of the list where the weight is important” [47].

Several computational studies to repurpose drugs have used BEDROC as a metric, albeit not to evaluate drug repurposing performance. Specifically, Govindaraj et al. use BEDROC to assess the ability of their algorithm to detect protein pockets binding similar ligands [74]. Alberca et al. use BEDROC to assess virtual screening of protein-ligand interactions [75]. Ajay Jain reports on limitations of BEDROC and other metrics that assess early enrichment from a virtual screening perspective, stating that they are biased to report elevated values based on the total number of positives and negatives [76]. An example of explicit use of BEDROC in drug repurposing is from Arany et al [77], who use it (along with AUROC) to evaluate effectiveness of their methods to produce drug rankings with respect to correct Anatomical Therapeutic Chemical (ATC) codes [77].

Comparing BEDROC scores across different *α* values is not advised [47]. For a given drug repurposing technology, users may seek to predict novel drugs across a multitude of indications with highly variable numbers of associated drugs. If an indication has 200 associated drugs, then *α* should be low to maintain the ≪ 1 condition. However, such an *α* may inappropriately lead to highly variable BEDROC scores for indications with a low number of drugs, as small changes in absolute ranking of these drugs correspond to a large change in the relative ranking. These requirements of *α* possibly lower cross-platform comparability; regardless, BEDROC is best applied to evaluate repurposing technologies using similar quality and size data (i.e., similar numbers and quality of drugs, indications, and corresponding associations).

Figure 5 illustrates the results of integrating BEDROC into CANDO. Most uses of BEDROC are based on single *α* value across all known drug-indication associations for all calculations to maintain comparability. As platforms become larger and more diverse, finding a balance to handle indications with varying number of associated drugs will be necessary, without being singularly biased to a few well-studied indications. Regardless, BEDROC has several major improvements over AUROC, and the former should be preferred when reporting results of drug repurposing technologies.

**Figure 5:**
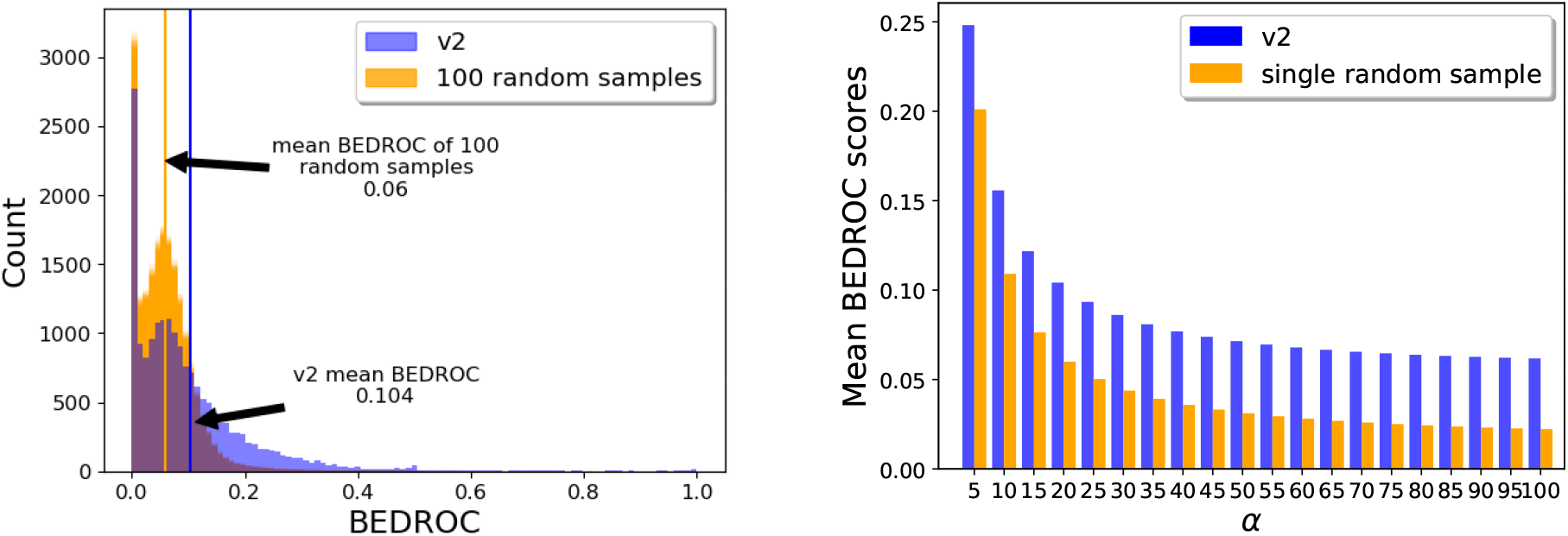
Evaluating CANDO performance using BEDROC. The left panel illustrates the count of CANDO v2 BEDROC scores using *α* = 20 for all compound-indication pairs in CANDO, compared to 100 samples obtained using shuffled CANDO data (orange). Using *α* = 20, the most commonly used value in the literature, the mean v2 BEDROC score is 0.104, compared to an average of 0.06 over 100 random samples, indicating that v2 outperforms this random control at retrieving known drug-indication associations on average. Generally, BEDROC scores for predicted drug-indication associations from v2 are considerably better than random, indicating their greater real world utility. The right panel shows the mean v2 BEDROC scores (blue) compared to a single random sample (orange) using *α* = 5, 10, …, 100. The random sample was constructed via shuffling of v2 data to obtain drug-drug similarities expected by chance, as usual. At higher values of *α*, certain drug-indication pairs may violate the conditions necessary for BEDROC to remain useful.

### 2.9 Discounted Cumulative Gain and Normalized Discounted Cumulative Gain

The discounted cumulative gain (DCG) is constructed with assumptions that top ranked results are more likely to be of interest, and that particularly relevant results are more useful [78]. While data in the form of ranking are readily measured by the DCG, classification schema should be converted to a ranking underpinning the decision boundary to be appropriately measured by DCG. The Ideal DCG (IDCG) is the DCG calculated for a ranking where all known actives are ranked the very best in the prediction list. The Normalized DCG (NDCG), with a value between 0-1, is obtained by dividing the DCG by the IDCG. The NDCG enables comparison and contrasting of performance evaluation with different numbers of relevant results with meaningful interpretation, i.e., we can use a single value to determine goodness of a drug-indication ranking/classification with greater confidence than most other metrics even when there are different numbers of associations.

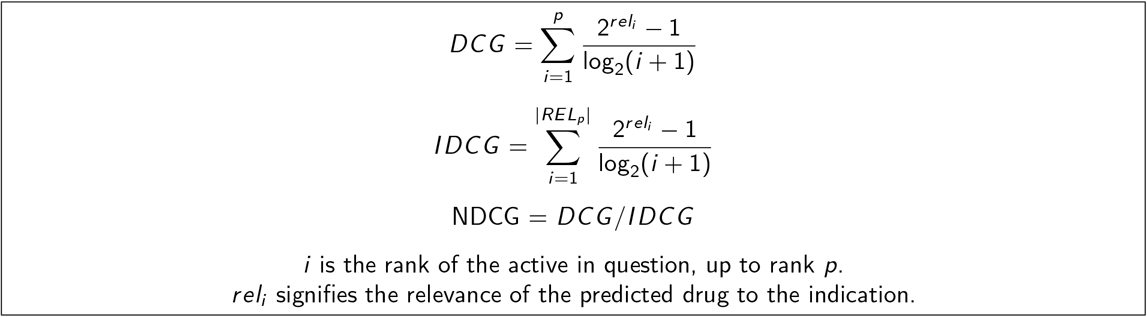

The value of *p* is a specific position (ranking) at which the NDCG is calculated. Therefore results are reported as NDCG_p_. Wang et al. suggest selecting *p* as a function of the size of the libraries used [79]. The distribution and measures of central tendency of NDCG at a cutoff or multiple cutoffs can be reported, i.e., a NDCG value can be calculated for every possible ranking. One of the most appealing features of DCG is the ability to assign relative importance, captured in the *rel_i_*, value. For a certain indication, there may exist more or less effective therapies, which can be reflected in the drug-indication association standard. Applied to precision medicine, such a relevance could hypothetically be determined on a per patient basis. This is a complex variable with a range of possible values, but is often used in a binary fashion. The NDCG is the best measure of correctness if a standard has known relative importance assignments.

Ye et al. use NDCG as the metric of choice for analyzing repurposing opportunities based on drug side effects [80]. Specifically, the authors report the top 10 ATC therapeutic categories with NDCG_5_ [80]. Specifically, the authors report the therapeutic categories of drugs (as classified by ATC codes) which achieved an NDCG_5_ of more than 0.7 during benchmarking. While the metric enables comparability, predicting a general categorization of a drug-indication association is easier than a very specific therapeutic prediction. In addition to using NDCG, Ye et al. report on putative therapeutic predictions that overlap with previous similar work [81]. Saberian et al. use NDCG to assess performance of their drug repurposing method for three indications, breast cancer, idiopathic pulmonary fibrosis (IPF), and rheumatoid arthritis (RA), with the score based on the rank of the left-out drug in each round of sampling [82]. Their work includes a sample calculation of NDCG in the corresponding Supplementary Material, compared to results based on 1,000 random rankings of drugs, though the mean NDCG of these rankings is not reported [82].

Figure 6 illustrates the mean NDCG at all cutoffs for CANDO v2. We obtain a mean NDCG_10_ of 0.060, compared to an average of 100 random data sets of 0.0197, and a theoretical average of 0.0199. Comparing these values along with the results shown in Figure 6 indicates that using CANDO to predict drug-indication associations has utility. The elevated cross-platform comparability of NDCG due to the use of logarithmic scoring and normalization makes it among the most useful metrics reviewed to measure success when used in drug repurposing technologies (Figure 7).

**Figure 6:**
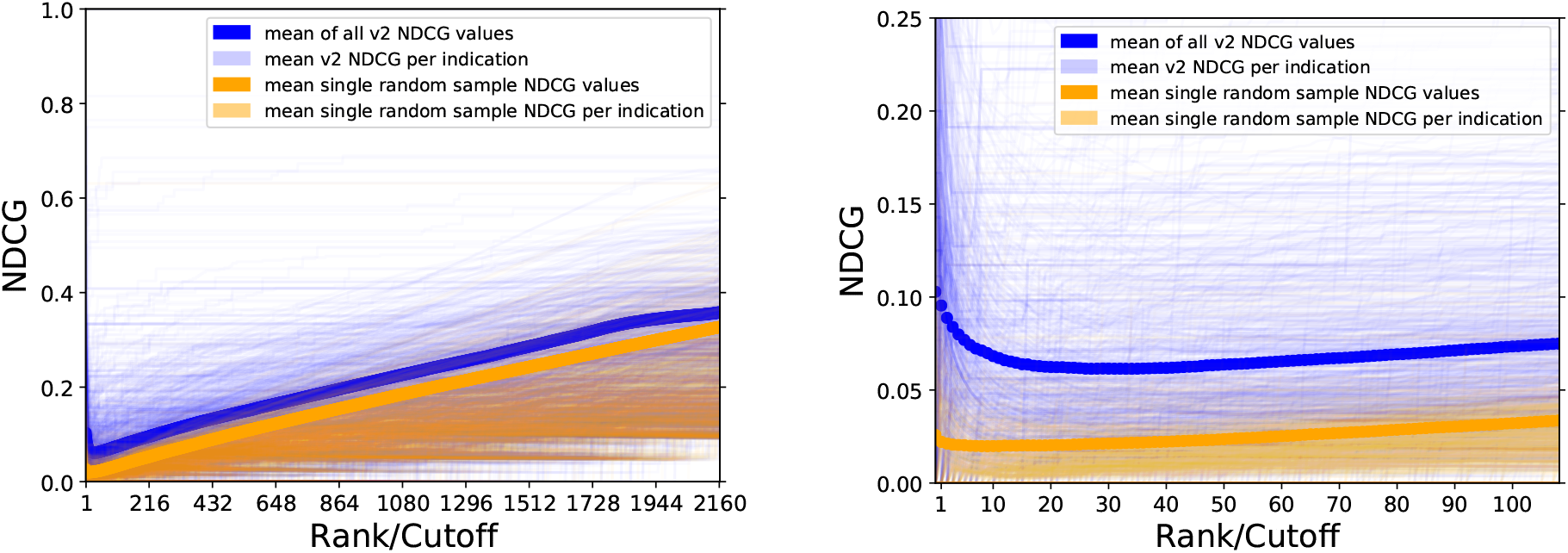
Evaluating CANDO performance using NDCG. The NDCG scores of CANDO v2 compared to those from a single random control at all ranks/cutoffs is shown on the left and at the top 20% of ranks (432) on the right. The overall mean of v2 is the dark blue line, with per indication means in light blue. The mean of a single random sample is in red, and the per indication means of the same sample is in orange. By chance, some predictions will be worse than random, as is evident whenever an orange line is higher than a light blue line but on average, v2 performs better than random at all ranks. These comparisons indicate the utility of CANDO at predicting drug-indication associations using the most rigorous performance evaluation metric considered in this study.

**Figure 7:**
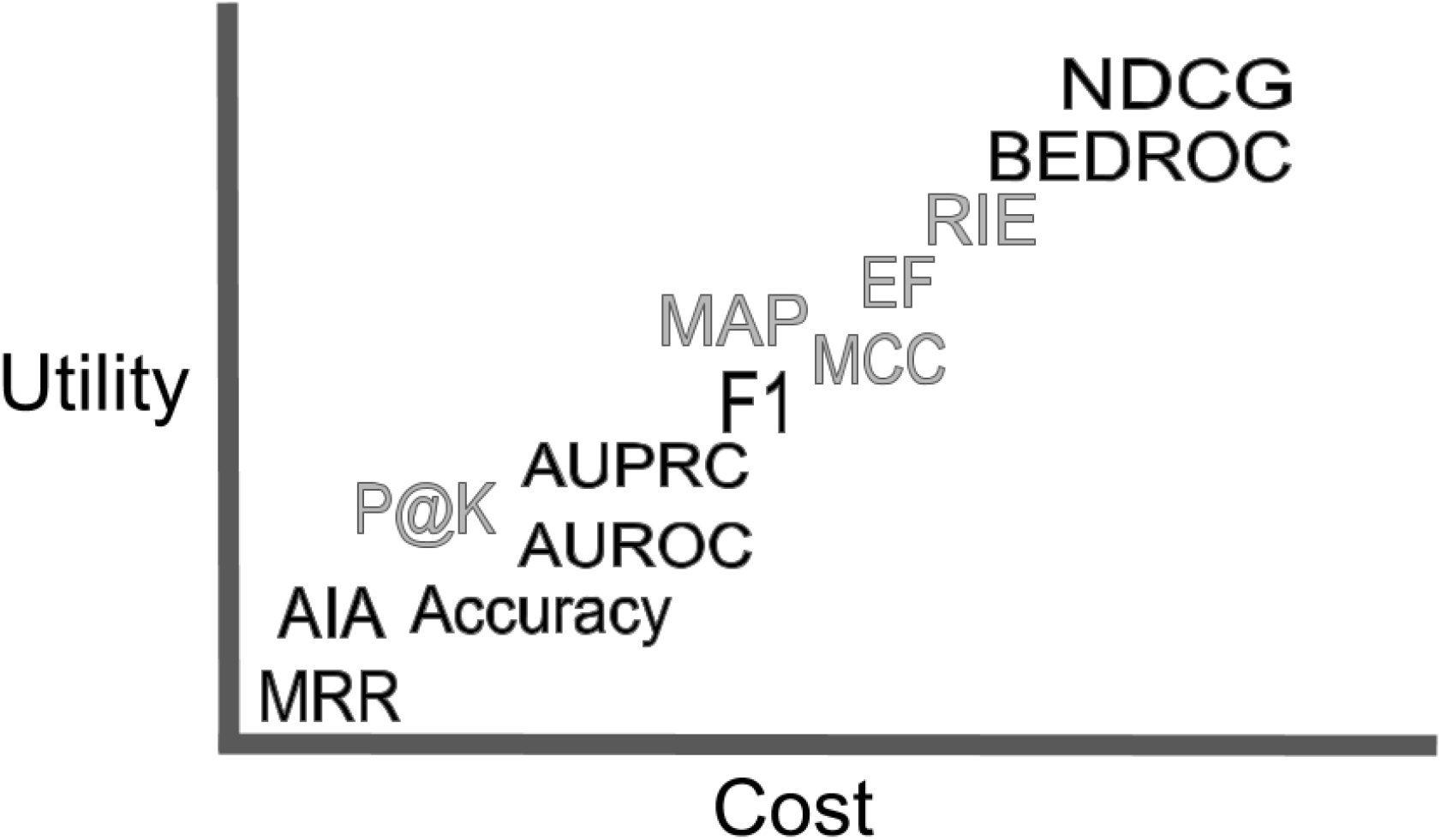
A subjective illustration of the relationship between cost and utility of performance evaluation metrics for drug repurposing. Further right on the horizontal cost axis equates to technologies requiring increasing computational power with decreasing intuition. The vertical utility axis is a gauge of enabling cross-platform comparability and, ideally, fidelity to reality through likelihood of success in prospective external validations for multiple drugs per indication. Metrics discussed in the Supplementary Material are in grey. We encourage scientists to use metrics such as NDCG and BEDROC, given their high utility and low incremental cost, in addition to avoiding overtraining using a single metric. This study makes the argument for the same general trend to be borne out through prospective clinical validation experiments of drug repurposing technologies.

### 2.10 Custom methods of performance evaluation

We have used our AIA metric in CANDO extensively [7–17] (section “Drug/compound characterization, benchmarking, evaluation metrics, and performance”). Similarly, Peyvandipour et al. used a custom evaluation metric in their systematic drug repurposing study [83]. The goal of this review and study is to more readily compare our results with others, an outcome toward which we are continuously striving, presently by integrating more widely used metrics into our system, and advocating for others to do the same. We initially developed AIA in response to our validation partners seeking a singular successful hit for an indication they were studying; similarly, other researchers may desire to use their own evaluation methods for their own reasons. Nonetheless, we recommend researchers also report results using one or more of the metrics described herein.

### 2.11 Evaluating drug repurposing in the context of precision medicine

A specific medication may be more or less efficacious for a particular person with a specific disease at a given time. Drug repurposing technologies can be tailored to arbitrary individual contexts and so precision/personalized medicine are growing areas of interest [32,59, 84–86]. Our group is currently exploring opportunities in the realm of precision cancer therapeutics using both CANDO and our molecular docking protocol CANDOCK [16]. We have previously published studies to predict HIV drug regimens based on the viral mutations circulating within a patient [87–89], understanding polymorphisms in the malarial parasite *P. falciparum* [?, ?, ?], explaining warfarin resistance [?], among many others. All these studies would have benefited tremendously from the application of evaluation metrics described in this study.

In the case of rare disease, including rare genetic diseases and rare cancers [32,90–92], computational drug repurposing experiments may offer the best chance at discovering efficacious treatments [3]. The field has promising initial results [84, 85, 93], but notions of correctness remains limited to mechanistic understanding and preclinical corroboration [32]. The use of particular metrics and quantifiable comparison between experiments is unknown, as it has not been done, but the metrics reviewed herein may have the same utility as when used broadly. Due to the low number of individuals with rare diseases, clinical trials are difficult to conduct, and only the most scientifically rigorous preclinical predictions with greatest confidence from drug repurposing technologies should be considered for further downstream research and use [90].

## 3 Cross platform comparability and fidelity to reality

### 3.1 Different standards, imbalanced data, suboptimal design

CANDO was designed to be a shotgun repurposing technology, i.e., to generate putative drug candidates for any/every indication. Lack of consensus with respect of performance evaluation makes it difficult to assess if this is true of any drug repurposing technology. A key component of drug repurposing technologies is the use of some standard drug-indication association library to which results are compared. A “perfect” evaluation metric cannot correct for data that does not reflect reality, is imbalanced, is over represented by “me too” drugs/compounds, or generated as a result of poor cross validation or over training.

There are a limited number of high quality data sources available for curating drug repurposing standards, and the ones that do exist fluctuate significantly. As an example, consider the drug ofloxacin, which is commonly classified as a fluoroquinolone antibiotic. Different standards associate it between one to seventy indications [39, 45, 94–96], which subsequently influences the odds of making a correct prediction by chance. Ultimately, different standards make it difficult to compare technologies and platforms.

Moving forward, differences in drug-indication association standards may be overcome through increasing consistency of drug classes across standards [97], or through integration of ontological understanding into all aspects of drug repurposing [98], using ontologies specifically designed for this purpose [99]. Knowing the ground truth is a prerequisite for measuring performance [100], and the use of scientifically rigorous ontologies will ensure robust modeling of reality.

It is natural there are different numbers of drugs associated with each indication within a particular single standard used in a large platform such as CANDO due to biological, economic, and even political reasons. The result is a large discrepancy in the number of drugs approved for or associated with a particular indication. For example, in the CANDO v2 drug-indication association standard, there are 218, 216, and 207 drugs associated with pain (MeSH ID: D010146), hypertension (MeSH ID: D006973), and seizures (MeSH ID: D012640), respectively, versus 8 for dermatomyositis (MeSH ID: D003882). While the performance of a system such as CANDO averaged over many indications is quite robust, performance on a smaller basis is more variable. Baker et al. identify hyperprolific drugs which have been studied in the context of many indications, and indications for which many drugs have been investigated as a treatment [101]. In an attempt to partly overcome variation in results and performance due to chance, Zhang et al. eliminated indications with less than ten associated drugs and drugs with less than ten associated indications from their platform [102]. Unfortunately, this action seems to contradict their attempt to meaningfully compare to the PREDICT project using the AUROC, as the distribution of drug-indication associations in PREDICT was greatly enriched in an opposite manner for drugs with less than ten indication associations, and indications with less than ten drug associations [60].

In describing the evolution of CANDO we use measures of performance evaluation applied globally, as we have done here. We also apply these measures to specific individual indications, particularly with respect to prospective validation of the platform or its components [7,8,10,17,89,103–105]. Regarding any classifier technologies, David Hang states, “[A] potential user is not really interested in some ‘average performance’ over distinct types of data, but really wants to know what will be good for his or her problem, and different people have different problems, with data arising from different domains. A given method may be very poor on most kinds of data, but very good for certain problems” [106]. In particular, it is easy to use different input data or comparison standards to obtain numerically better results. Since a great average performance overall does not guarantee similar performance for specific drugs/indications, and vice versa, drug repurposing technologies must undergo a thorough vetting across multiple libraries, standards, and experiments (indications to which the technology is applied) to be considered robust.

### 3.2 Inherent limitation of claims and metrics

Metrics for evaluating success of drug repurposing typically rely on the assumption that all associations not part of a standard are negatives. This goes against the entire premise of drug repurposing, which is to expand the list of known drug-indication associations, and any prediction made that is not present in a standard could subsequently be proved correct.

A perfect score for a drug repurposing experiment based on some evaluation metric does not necessarily mean that perfect drug repurposing success was achieved. Numerical evaluation is limited by the choice of cutoff. If the results are an ideal ranking or classification of drugs that are known to be a treatment for an indication against those that are not, then all of the metrics discussed here will yield their best possible result. The actionable information, however, is in those drug-indication associations which are truly novel and even unexpected. For example, in a drug repurposing experiment for breast cancer, the top ten putative therapeutics reported are known drugs to treat this indication, and the first novel prediction is at rank eleven [107]. If any of metrics described here was used for evaluation with a rank/cutoff of top ten, then this experiment would achieve a perfect score without discovering a novel drug to repurpose.

A purported benefit of high-throughput approaches are the vast size and quick speed enabling us to explore and make discoveries more quickly than ever before. Because of this, specific elements of data/standards used in drug repurposing experiments and qualitative results (predictions) may sometimes be clinically wrong or nonsensical. This includes incorrect notions of indications [108] and proposition of treatments known to exacerbate disease [109,110]. A small number of mistakes or inconsistencies in a large drug repurposing technology do not necessarily invalidate it, but necessitate the need for manual expert inspection and curation.

As artificial intelligence (AI) and machine learning become more prevalent for drug repurposing [73], biased data and overtraining may return results that are falsely interpreted as being significant or high confidence. Mindset, culture, and willingness to apply computational drug repurposing models and use their results, considered relevant for the success of AI in drug discovery and development [111], will partly depend on the confidence in the methods and output, as evidenced by how we evaluate performance. Orthogonal metrics which capture different aspects of the goodness of an experiment could be used in concert to overcome bias and potential voodoo science pitfalls. In the future, prospective blinded assessment of computational drug repurposing, such as those used in protein structure prediction and molecular docking [112,113], may be another solution to alleviate the problem of bias.

Complex yet quick drug repurposing technologies can render results beyond the cognitive ability of a person to be familiar with all of its components and the amount of resulting data. The use of rigorous performance evaluation metrics enable a culture where scientific rigor and correctness is valued more than the novelty in making claims of putative repurposed therapeutics. Even so, it would be prudent for basic science researchers to work with clinicians to ensure that their results make sense in order to guide correct predictions into clinical use and improve human health.

### 3.3 Validation

In the context of drug repurposing technologies, “validation” may refer to: internal validation through testing of models on unknown or hidden data; performance as evaluated by the types of metrics discussed here; or to some external independent corroboration. The latter form of validation may refer to anything from selective reporting of similar results in the literature to results from prospective preclinical and/or clinical studies.

A popular strategy for drug repurposing is to report corroboration of predictions made using computational methods with previously reported independent research in the literature, in a case-based or large-scale analysis [18, 52, 114–116]. We have used this strategy [10, 11, 17], including highlighting literature that contradicts our findings [14]. It is relatively easy with this approach to find examples which support preformed conclusions, and report only those, representing potentially serious instances of confirmation bias. Selective literature corroboration is neither systematic nor hypothesis-driven.

Similar to literature analysis is using data on clinical trials completed or in progress [66, 72, 82, 117, 118]. Through analysis of electronic health records, preventative associations between drugs and indications, i.e., form of drug repurposing, have been discovered in an ad hoc manner [119, 120].

There are several examples of preclincal (*in vitro*, *in vivo*) validation done following a computational drug repurposing experiment [18, 20, 59]. In the future, some of these technologies may approach or even rival the current best method for elucidating the usefulness of a drug, which for now remains double blind, placebo controlled, randomized trials with clinically relevant primary endpoints (prolonged life or improved quality of life), and representative samples of subjects, to evaluate both efficacy and safety [121].

The goal of achieving successful drug repurposing, from technology to clinic (Figure 1)), is still mostly aspirational at this stage. However, progress is being made. The most successful discoveries made using drug repurposing technologies are ad hoc singular events which current metrics are not well suited to evaluate [122,123]. While emotionally unsatisfying, we can search for and use metrics which enable us to compare our technologies directly to each other, for the sake of rigorous science, intellectual merit, and broader impact.

### 3.4 Response to pandemics and novel disease

The potential for drug repurposing technologies to help respond to epidemics and pandemics rapidly, side stepping lengthy, costly preclinical and clinical studies, is enormous. Recent examples include the Ebola virus disease West African outbreak of 2014, the emergence of the Zika virus, and the COVID-19 global pandemic caused by the novel coronavirus SARS-CoV-2. In all instances, there were no drugs approved to treat these indications, but drug repurposing technologies were used to generate putative therapeutics quickly [10,17,124–129]. In these examples, it is challenging to use the metrics we have described to evaluate the predictions as there are no previously approved treatments. However, if a platform or methodology has reported measurements of success, especially in related indications to prevent, treat, or cure viral infections, then relying on those performance values as a quantified surrogate may have utility.

The COVID-19 pandemic also illustrates a potential downside of quickly available drug repurposing predictions. Drugs that have been predicted to be efficacious in treating an indication may have serious side effect profiles, or unknown side effects when used in different quantities to treat novel indications. Studies of rigorous evaluation of drug repurposing platforms expressed in clear and precise language will help scientists, healthcare workers, institutional and government officials, and the public make informed judgements with respect to future steps on how to use the generated drug candidates for a given indication.

## 4 Conclusion

Drug repurposing will help advance and evolve therapeutic discovery in the 21st century, bringing new medicines to patients in need. Advancing the field depends on whether we can rigorously evaluate the validity and meaning of our computational repurposing experiments with confidence, a critical component of platform development. We have shown how integration of disparate metrics into the CANDO platform supports this claim. The metrics currently used for gauging correctness of drug repurposing technologies vary in terms of enabling cross-platform comparability, as well as eventual clinical use of predicted therapeutics. The development and use of improved evaluation metrics will enhance cross-technology comparability and enable more accurate modeling of reality to deliver on the potential of drug repurposing.

## 5 Acknowledgements

The authors thank Dr. Manoj Mammen and other members of the Department of Biomedical Informatics at the University at Buffalo for critical reading of this manuscript. This work was supported in part by a National Institute of Health Directors Pioneer Award (DP1OD006779), a National Institute of Health Clinical and Translational Sciences (NCATS) Award (UL1TR001412), NCATS ASPIRE Design Challenge Awards, a National Library of Medicine T15 Award (T15LM012495), a National Cancer Institute/Veterans Affairs Big Data-Scientist Training Enhancement Program Fellowship in Big Data Sciences, and startup funds from the Department of Biomedical Informatics at the University at Buffalo.

## Notes

### Competing Interest Statement

The authors have declared no competing interest.

