## Supplementary Information for "Evaluating Performance of Drug Repurposing Technologies"

### 1 Additional metrics of interest

The metrics described are important for historical and developmental reasons. They are often used as part of a drug repurposing experiment, but are not used to describe the putative drug-indication association results.

#### 1.1 Matthews correlation coefficient

The Matthews Correlation Coefficient (MCC) is one metric which summarizes a confusion matrix (all TP, FP, TN, FN) into a single value. It can be calculated using the formula [1]:

$$MCC = \frac{TP \times TN - FP \times FN}{\sqrt{(TP + FP)(TP + FN)(TN + FP)(TN + FN)}}$$

Applicable to many tasks of binary classification, Chicco and Jurman provide a careful analysis of the MCC and conclude, "[MCC] should be preferred to accuracy and F1 score in evaluating binary classification tasks by all scientific communities" [2]. We have yet to find an example of MCC used in a ranking scheme of drug repurposing. Yang and Argawal use the MCC as a metric for evaluating correctness of drug-side effect associations, but used AUC when evaluating drug-indication associations [3]. Donnet et al. use the MCC as a metric for evaluating performance of their system to rank drugs by similarity to a query drug [4]. The authors point out what a perfect performance based on internal evaluation of their system would be. Li et al. combine information of molecular structures and clinical symptoms using deep convolutional neural networks to capture information of drug-indication associations, and report results using MCC among several other metrics [5]. While not a drug repurposing paper per se, as they are not suggesting novel uses for approved drugs, Russo et al. focus on virtual screening against a particular target [6]. The authors use a number of different metrics in evaluating performance of their deep learning models: recall, precision, F1-score, accuracy, AUROC, Cohen's Kappa, and Matthews correlation coefficient, allowing the reader to gauge relationships between metrics. Moreover, results are illustrated using figures that facilitate easy comparison [6]. We report the mean MCC value of 0.01817 over all drug-indication pairs in CANDO v2 at the top 100 cutoff, compared to -0.00029 obtained from an average of 100 random runs ($\approx 0$, as expected).

#### 1.2 Precision-at-K, Average Precision, Mean Average Precision

When reporting the precision of ranking for drug repurposing, the specific rank at which that measurement was calculated must be reported. Due to the nature of a ranking scheme, precision by definition refers to precision-at-K (P@K), where K is the rank. Computational drug repurposing to associate drugs with indications is analogous to information retrieval and web web search, and hence use of the precision-at-K metric is apt [7]. Kuang et al.

report precision at a K of 8, 16, 24, 32, and 40. Explicitly reporting the average precision is better than reporting the mean precision at a given cutoff across an arbitrary set of measurements. The mean precision of a drug repurposing experiment is reported by Xu and Wang [8], and Bakal et al. [9]. At the top 40 rank/K, CANDO v2 obtains a mean precision-at-40 of 0.034, compared to 0.017 of 100 random runs using the same pipeline.

Average precision as defined from an information retrieval perspective is the mean precision-at-K for each K of a relevant item [10]. Precision at these specific ranks are a subset of precision at all ranks.

$$\text{Average precision} = \frac{1}{N}\sum_{K} P@K$$

where $K$ is the rank of each known active, and $N$ is the total number of compounds

Because average precision concerns relevant items, the value $K$ is dependent upon the standard used. In contrast, with $precision - at - K$, $K$ is any value chosen by the investigator. Re and Valentini use the average precision when "prioritizing drugs in integrated biochemical networks according to specific DrugBank therapeutic categories" [11]. Mean average precision (MAP) is the mean of all average precision values. The MAP was used by Yang et al. and reported on a top percentage basis about their experiments repurposing drugs [12]. For experiments reporting a single ranking of drugs predicted to treat a single disease, calculating a mean average precision is not possible.

### 1.3 Enrichment Factor

Enrichment Factor (EF) is "the measure of how many more actives we find within a defined 'early recognition' fraction of the ordered list relative to a random distribution" [13]. When EF is used in a drug repurposing technologies, it is most often used as an intermediate measurement, e.g., assessing how well a platform is able to capture known drug-target data as part of a larger experiment where such data is used to predict novel uses for known drugs. We have yet to find an example of EF being used to directly measure correctness of predicted drug-indication associations.

EF is often reported as $EF@n$, where $n$ is the number of results considered, i.e., calculate the EF considering the top $n$ drugs/compounds. Historically, common values for $n$ include 10, 20, 50, or 100. In CANDO it is easy to calculate EF at all $n$, i.e., at all possible cutoffs. This helps foster comparability between platforms which use similar library sizes.

EF lacks discrimination power within a particular cutoff. For example, 10 actives ranked the very best within the top 100 compared to 10 actives ranked the worst within the top 100 would score the same [13]. Additionally, EF is dependent on the number of compounds and active compounds [14], which limits interpretation of numerical results and comparability between platforms. As EF is non-scaled, it is difficult to interpret visualized results, as seen in Figure 1. Numerically, using CANDO v2, the mean EF value over all drug-indication pairs at the top 100 cutoff is 2.02, compared to 0.98 obtained from 100 random samples.

### 1.4 Robust Initial Enhancement

To date, we have yet to find an example of a "pure" drug repurposing experiment using robust initial enhancement (RIE) as a main metric, but instead have found RIE to be used when describing results of virtual screening and target prediction of compounds [15]. BEDROC is a score derived from RIE, with less shortcomings, namely lack of normalization [13]. Thus details about RIE apply to the more widely used BEDROC (described in the main text), so we recommend using the latter. For use in describing BEDROC, we will define RIE here, using the notation of Truchon and Bayly, mentioning $x_i$ is the relative rank of an active, i.e., $\frac{r_i}{N}$, where $r_i$ is the rank of the active, N is the number of compounds in the set, n is the number of actives, and $\alpha$ is a parameter at the choice of the investigator (see description of BEDROC in the corresponding manuscript).

$$RIE = \frac{\frac{1}{n}\sum_{i=1}^{n} e^{-\alpha x_i}}{\frac{1}{N}\left(\frac{1-e^{-\alpha}}{e^{\frac{\alpha}{N}}-1}\right)}$$

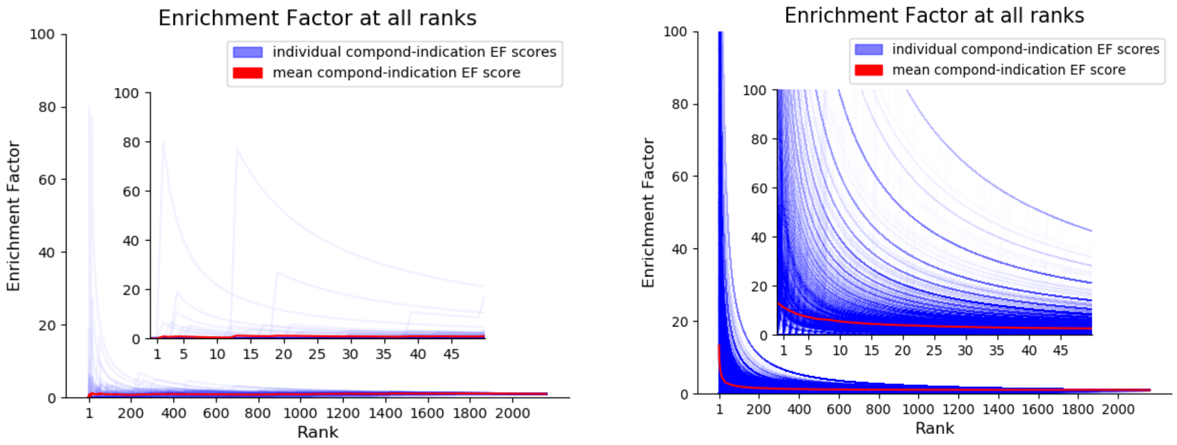


(a) (b)

Figure 1: **Evaluating CANDO performance using Enrichment Factor.** The Enrichment Factor (EF; vertical axis) for a random subset of 100 drug-indication pairs at every rank (horizontal axis) in CANDO v2 (blue) is shown. The EF decreases as the rank increases. The average EF for that set of drug-indication pairs at every rank is shown in red. In a platform such as CANDO with thousands of drug-indication pairs, it is difficult to discern meaning of the visualization of this non-normalized data. The left panel (a) shows a random subset of drug-indication pairs at all cutoffs, whereas the right panel (b) shows the Enrichment Factor for all pairs.
